# Anglers as Potential Vectors of Aquatic Invasive Species: Linking Inland Water Bodies in the Great Lakes Region of the US

**DOI:** 10.1101/2022.09.29.510070

**Authors:** Stephen J. Morreale, T. Bruce Lauber, Richard C. Stedman

**Author notes:** Corresponding author (SJM). These authors contributed equally to this work.

## Abstract

Unimpeded transfer and spread of invasive species throughout freshwater systems is of global concern, altering species compositions, disrupting ecosystem processes, and diverting economic resources. The magnitude and complexity of the problem is amplified by the global connectedness of human movements and the multiple modes of inter-basin transport of aquatic invasive species. Our objective was to trace the fishing behavior of anglers delineating potential pathways of transfer of invasive species throughout the vast inland waters of the Great Lakes of North America, which contain more than 21% of the world’s surface freshwater and are among the most highly invaded aquatic ecosystems in the world. Combining a comprehensive survey and a spatial analysis of the movements of thousands of anglers in 12 states within the US portion of the Great Lakes Basin and the Upper Mississippi and Ohio River Basins, we estimated that 6.5 million licensed anglers in the study area embarked on an average of 30 fishing trips over the course of the year, and 70% of the individuals fished in more than one county. Geospatial linkages showed direct connections made by individuals traveling between 99% of the 894 counties where fishing occurred, and between 61 of the 66 sub-watersheds in a year. Estimated numbers of fishing trips to individual counties ranged from 1199–1.95 million; generally highest in counties bordering the Great Lakes. Of these, 79 had more than 10,000 estimated fishing trips originating from anglers living in other counties. Although angler movements are one mechanism of invasive species transfer, there likely is a high cumulative probability of invasive species transport by several million people fishing each year throughout this extensive freshwater network. A comprehensive georeferenced survey, coupled with a spatial analysis of fishing destinations, provides a potentially powerful tool to track, predict, curtail and control the transfer and proliferation of invasive species in freshwater.

## Introduction

Vast amounts of effort have been dedicated to detecting, understanding, controlling, and predicting the spread of invasive aquatic plants and animals. The resources expended have only hinted at the magnitude and scope of the problem, with estimates exceeding 14,000 alien species in Europe [1] and more than 50,000 in North America, and with the costs of managing invasives in those regions easily exceeding $100 billion (USD) annually [2–4].

Over recent decades, increased global connectivity has resulted in the spread of invasive aquatic species throughout both marine and freshwater systems. In North America, freshwater invasive species include algae such as didymo; a host of other aquatic plants such, as *Hydrilla* and water chestnut; and animal species, ranging in size and diversity from spiny water fleas and New Zealand mud snails, to Asian carp and nutria. Transmission and spread of aquatic invasives in freshwater can occur through multiple means, including extreme storm events [5], natural dispersal and range expansion, intentional stocking, direct connections through canals and inter-basin transfer of water, transport by shipping [6], and behaviors associated with tourism and recreation (for a comprehensive review, see [7])

The wide variety of invasive aquatic organisms; the numerous possible vectors and modes of transport and transmission; and their ubiquity, from the smallest to the largest water bodies, all contribute to an extensive array of unique characteristics that make each freshwater invasion event different. Accordingly, there is no apparent single method of detection, prediction, or control of transmission. There have been numerous and varied approaches to detection, including basic manual sampling and monitoring [8] and inspections [9]; stakeholder and participant surveys to gauge presence and magnitude of impact [4,10] or develop conceptual models [11]; and analyzing eDNA [12]. Different approaches focus on modeling and predicting spread using risk assessment and screening tools [13,14], predicting egg transport [15], modeling propagule pressure [6,16–18], and modeling of impedance surfaces to identify least-cost pathways of likely transmission [19]. Some of these techniques include gravity models, which commonly use empirical data on current source locations and likely destinations, based on proximity and connections through transportation pathways or human activities that increase risk of spread. Methods may also include forces of calculating the potential connections between water bodies of different sizes [20,21], or assessing the number of registered boaters [22] and the magnitude of recreational use [23,24], all of which can influence the likelihood of introduction and spread of aquatic invasives.

In the US, efforts aimed at curtailing the spread of invasives among freshwater bodies have disproportionately targeted commercial vessels relative to recreational boats. Commercial shipping is both concentrated (many fewer boats) and a more straightforward task to regulate: the release of commercial shipping ballast water is, at least superficially, a relatively simple task with a limited number of actors, known travel routes, and straightforward solutions [25,26]. This contrasts with recreational boats, which number in the millions in inland waters—and even more so with the far greater number of individual recreational anglers, who are widely scattered and whose extensive activities can represent an immense challenge to monitoring and predicting. The contribution of recreational boaters and anglers to the transfer of a wide range of aquatic invasive species between inland water bodies is important and poorly understood. For example, recreational boats readily move zebra mussels in water retained in live wells, entangled in plants, and attached to trailers, lines, and anchors [9,27,28]. Similarly, fragments of many macrophytes are easily transported by recreational boats introducing species into freshwater lakes [29]. Indeed, recreational boats and anglers are likely the primary vector for the transmission of exotic zooplankters and mollusks [30–33] into freshwater bodies. More generally, a wide range of aquatic and terrestrial organisms are carried between sites on boats and trailers, including macrophytes, zooplankton, mollusks, and aquatic insects [34].

The transport of aquatic invasives is not limited to transfer by boats, however. Individual recreational anglers play a key role in inter-basin transmission through release of bait and transport of equipment, such as buckets, nets, and even fishing line [4,35]. Although the potential impact of a single individual angler may be minimal, the sheer magnitude and ubiquity of movements of individuals between water bodies greatly increases the cumulative probability of transport and spread of invasive species by anglers. This also contributes to the immense difficulty of tracing movements of anglers and predicting and monitoring potential transmission pathways—especially compared to commercial shipping.

Despite their potential importance, attempts to assess the impacts of anglers on the spread of aquatic invasive species are rare in the literature, and tend to be embedded in studies on recreational boat movements [4,23,32,36–38] . A common obstacle to integrating movements of anglers into analyses and predictions of spread of aquatic invasive species is the sheer scale and interconnectedness of many freshwater systems. This is epitomized in the inland waters of the Great Lakes, which extend across an area of over 244,000 km^2^ in the Northern US and Southern Canada, and contain more than 21% of the world’s surface freshwater. In this system, aquatic invasive species permeate the vast waterway and its extensive network of tributaries, ranging from intermittent streams to some of the world’s largest rivers and lakes. The Great Lakes are among the most highly invaded aquatic ecosystems in the world, with an estimated new non-indigenous species discovered every 28 weeks in the recent past [26]. Further complicating attempts to predicting patterns of transfer of aquatic invasives are the extensive movements of recreational fishers in this vast system. For example, in a single year (2011) there were more than 6.6 million licensed anglers in the 12 US states comprising the Great Lakes and Upper Mississippi Basin; they fished a calculated total of 224 million days [39].

In our study, we tackled the large problem of attempting to trace the movements of anglers in this expansive US Great Lakes region. To complement previous granular and highly geographically focused studies, our objective in this study was to create a model that estimates patterns and magnitudes of annual movements of all licensed anglers throughout the US portion of the Great Lakes Basin and the Upper Mississippi and Ohio River Basins. Our research combined a comprehensive survey and a spatial analysis of the movements of anglers traveling to and from all the counties and sub-watersheds in the study region, encompassing 12 states within the northern and mid-western region of the US. It is our intent that mapped linkages between water bodies and the estimated magnitudes of movements of anglers along connecting pathways can be instrumental in focusing efforts to determine “when, where, and how to most effectively and cost-efficiently detect invasive species” [40], and can be used as a guide to facilitate sampling, prevention, and intervention strategies.

## Materials and Methods

### Study Area

Our study focused on anglers and their movements across 12 states in the northern and mid-western US bordering the Great Lakes and Upper Mississippi and Ohio River basins (Fig 1). The states included in the study were New York, Pennsylvania, Ohio, Indiana, Michigan, Illinois, Wisconsin, Minnesota, Iowa, Missouri, Kentucky, and West Virginia. The study region spanned an area exceeding 3.2 million km^2^ and included 1042 counties and 66 watersheds at the HU 4 hydrologic unit level [41].

**Fig 1.**
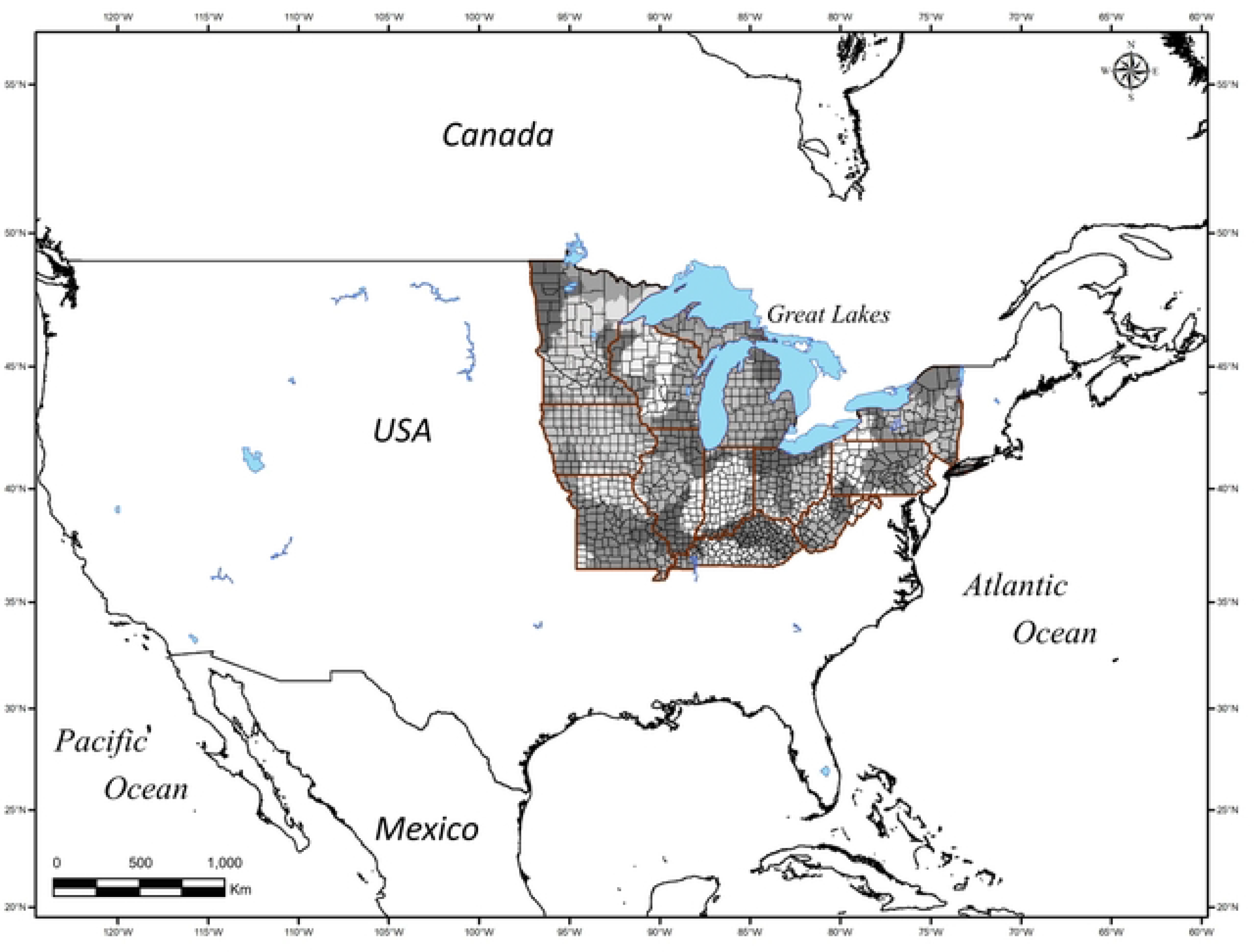
The study area encompassed 12 states 1042 counties and 66 watersheds (HUC4 Hydrologic Units) in the broader Great Lakes and upper Mississippi River basins in the northern and mid-western US. Surveys of 2576 licensed anglers who fished within the study area in 2011 provided a representative sample from which to estimate the behaviors and patterns of movements of more than 6.5 million licensed US anglers who fished in waters of those states in a single year.

### Angler Survey

In the spring of 2012, a survey was conducted of recreational anglers across the entire study area. The details of this survey were reported in previous reports [39,42]. The study population was defined as those individuals living in the 12-state region who were licensed to fish in 2011. The survey was conducted in two stages: (1) a screening survey conducted by telephone; and (2) a main survey conducted by mail or online.

First, a sample of anglers was recruited in each state through a screening survey in which eligible participants were randomly selected from fishing license records from 2011 in all survey states except Ohio and West Virginia (which would not release their fishing license data). Only anglers with addresses within the 12-state region were used in an initial sample of 28,200 fishing licenses in these 10 states, which was pared down by phone to a total of 7,201 individuals who met several screening criteria (to assess whether they fished in 2011 and intended to fish in 2012), and agreed to participate in a subsequent angler survey. Additional screening questions were designed to later aid in the assessment of non-response bias after the survey was complete. Licensed anglers from Ohio and West Virginia were recruited through random digit dialing [39]. Using the same screening process, nearly 17,000 telephone numbers were further pared down yielding 491 additional participants. These contributed to an overall total of 7,692 anglers that were recruited to participate in the survey of resident anglers across the 12-state study region. From the selected participants, data were collected through either a web-based survey (n=4,562), or a mail survey (n=3,112), based on participant preference. Non-respondents were sent reminder letters and an additional copy of the survey instrument. When survey respondents were compared to non-respondents to determine if non-response bias existed, the difference in intention to fish, was negligible, and there was no significant difference in the cumulative total of days fished [42]. Thus, we deemed that the behaviors of the survey respondents were largely representative of the fishing patterns of all licensed anglers in the 12 states.

Among the numerous survey questions in the broader study, those at the core of this spatial-analytical study included information on angler fishing behavior in 2011, including: 1) zip code of primary home, to provide a location of origin for fishing trips; 2) destination locations of all fishing trips taken in 2011, recorded at the county level; and 3) the number of trips taken to each fishing location throughout the year. Because we collected the location of the fishing trips at the county level, we eliminated those trips for which we did not have county data, which amounted to 5.3% of all trips, but resulted in the elimination of 27.1% of the respondents from the data set. Thus, the final data set used in our spatial analyses was based on 2576 individual respondents. In addition to estimating the overall fishing activity at each location, the fishing patterns, origins and destinations of all individual anglers across the Great Lakes region were spatially referenced and mapped to estimate the linkages and magnitudes of movement between and among separate counties and watersheds in a single year.

### Data Weighting

To scale up from the anglers represented in the survey, weights were estimated for anglers in each state based on the number of survey participants in proportion to the state’s registered fishing license holders for 2011—see [42]. Weighted values were used to estimate the total number of anglers who fished and the total number of individual fishing trips in all 1042 counties, and the magnitudes of inter-county linkages over the single year.

### Spatial datasets

Postal ZIP codes of survey respondents were spatially referenced by joining angler address data with the TIGER ZIP Code Tabulation Areas TIGER/Line Shapefile [43] To further anonymize participant addresses, ZIP codes were georeferenced to their respective counties [44], which were then listed as the origins of fishing trips.

Reported fishing destinations also were georeferenced using county boundaries. In addition, the 12-state study region is composed of all or part of 66 sub-watersheds that were spatially referenced using the USGS National Hydrography Dataset Best Resolution dataset (NHD) for Hydrologic Unit (HU) 4 [41]. Counties of origin and reported fishing destinations of all anglers in the survey were georeferenced and matched to their watershed. To establish a network of physical connections created by angler movements between counties, we mapped the linkages between counties whenever individual anglers fished in multiple locations in the single year. The same analytical process was used for delineating the movements of anglers and mapping their network of linkages among watersheds.

### Spatial Analyses

All analyses were conducted using SPSS, dBase PLUS 11, and ArcGIS 10.8. Data Layers for maps of angler fishing locations included counties (from cartographic boundary shapefiles), ZIP codes (from TIGER/Line shapefiles), and watersheds (from NHD HU 4 shapefiles). Codes for calculating weighted totals and inter-county and inter-watershed linkages were written in dBase Plus 11 and Python 2.7.18.

The magnitude of linkages between counties was calculated using spatially referenced fishing destination data from the surveys, which were recorded at the county level. First, to estimate the total linkages made by all anglers moving between different counties, we summed the weighted values for individuals that visited more than one county. We represented this visually by mapping and displaying the magnitude of the total connections between separate counties as a result of the movements of anglers in the course of the single year. These mapped connections can be used to determine the possible pathways by which invasive species may be transported throughout the freshwater network.

We also mapped out the network of connections made by individual anglers traveling between different watersheds. These linkages were calculated from the unweighted spatially referenced data of angler fishing behavior; depicted at a coarser resolution, by resampling to a watershed level (HU 4 hydrologic unit). All angler movement network maps were created using ArcGIS 10.8

## Results

Surveys of 2576 anglers that were issued a license and fished within the study area provided a representative sample from which to estimate the behaviors and patterns of movements of more than 6.5 million licensed US anglers who fished in those states in a single year.

### Fishing Trips

Respondents reported a combined total of 76,713 fishing trips, with an average of 30 fishing trips per angler over the course of the year. Fishing destinations were widespread across all 12 states and most of the counties. Only 148 (14%) of the 1042 counties in the study area were not reported to be fishing destinations by respondents for the survey year. Among the survey respondents, county totals ranged from 1 to 738 trips by anglers throughout the year. The calculated weighted sums of fishing trips to the counties (based on the number of licensed anglers in the states) ranged from 1199 to more than 1.95 million to a single county (Cook County, Illinois) in the single year of 2011 (Fig 2). Fishing destinations were generally heaviest in counties bounding the waters of the Great Lakes.

**Fig 2.**
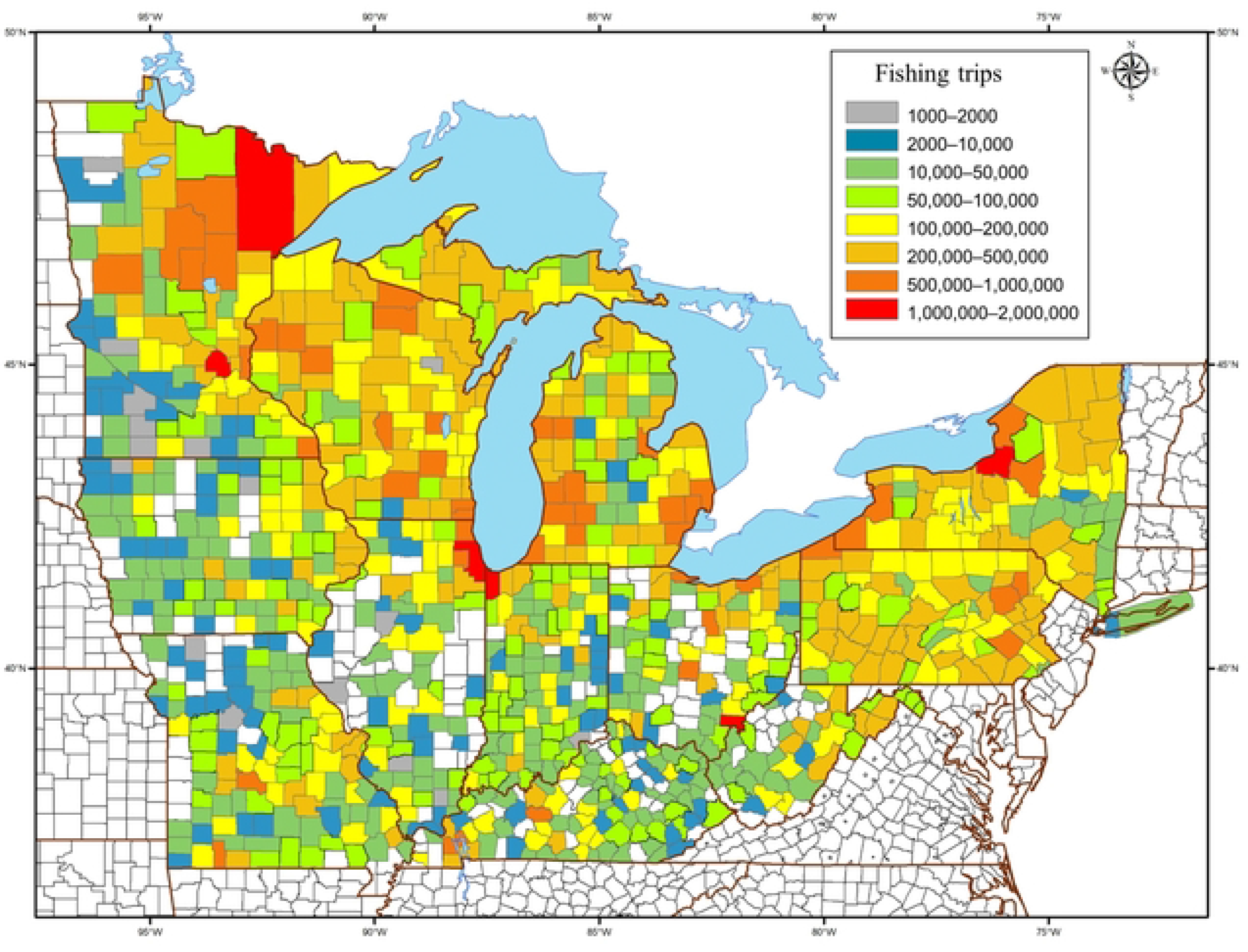
Weighted sum of all fishing trips by anglers in the broader Great Lakes region in a single year to counties in 12 US states surrounding the Great Lakes and upper Mississippi River basins. The estimated numbers of fishing trips with destinations to each of the 1042 individual counties ranged from 1199 (blue), to more than1.95 million (red) in the single year of 2011. Only 148 of the 1042 counties in the study area were not reported as fishing destinations (white).

### Fishing Trips Away from Home

By comparing fishing locations to addresses of all survey respondents, 94% of counties where fishing took place were reported as fishing destinations for anglers originating from another county. The calculated weighted sums for the year ranged from 1199 to more than 143,000 fishing trips to counties in the study area by anglers from other counties (Fig 3). Of the 85 US counties directly bordering the Great Lakes, 79 had more than 10,000 estimated fishing trips originating from anglers living in other counties in the survey area. The four counties with the highest numbers of fishing trips by anglers from a different county all were in Northern Minnesota. Only 51 of the 894 counties where fishing was reported had fishing trips that all originated within the same county.

**Fig 3.**
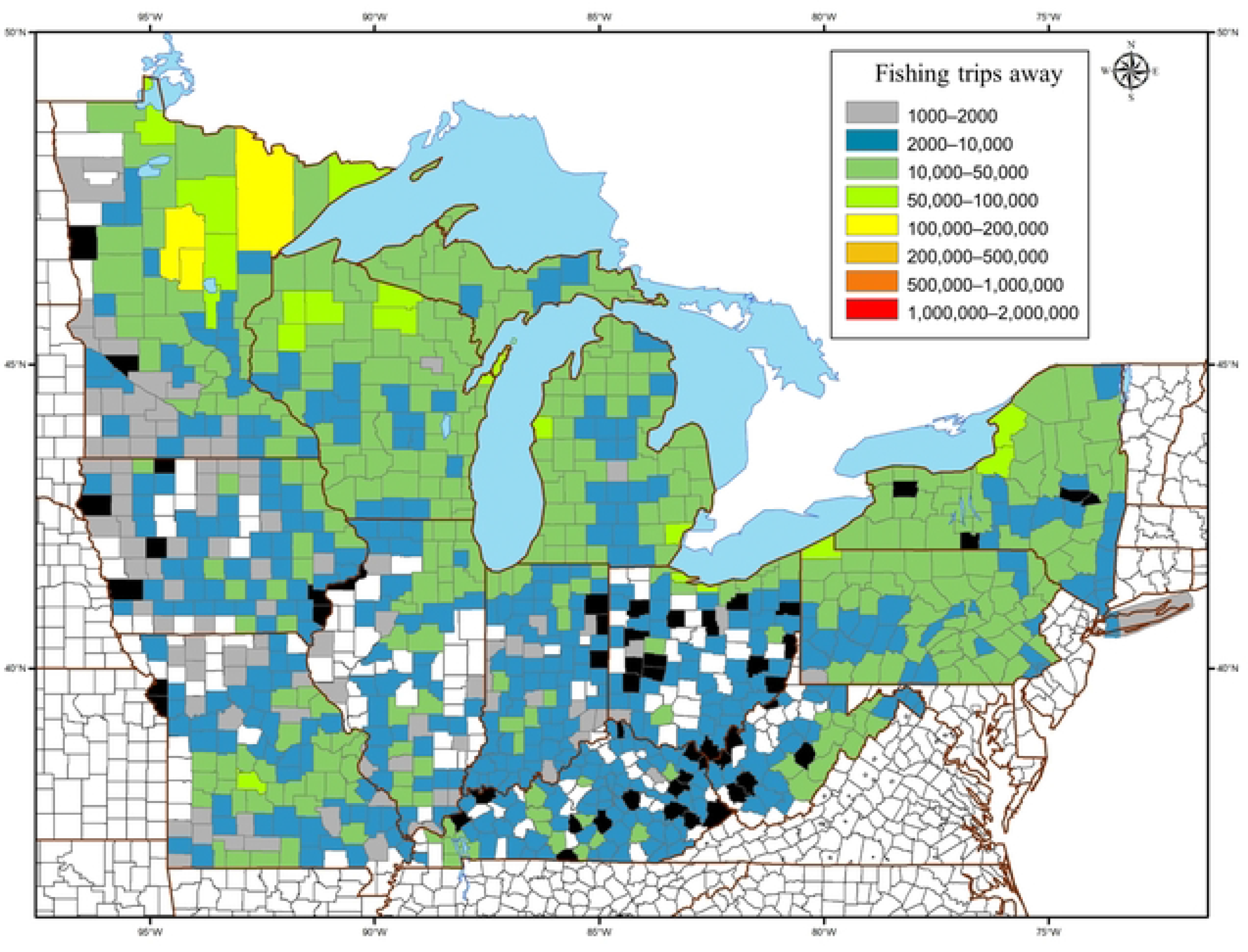
Weighted Sums of fishing trips to counties by anglers traveling from other counties in a single year in the broader Great Lakes Region. The calculated numbers of fishing trips to counties in the study area by anglers from other counties ranged from 1199 (blue), to more than 143,000 (red) in the single year of 2011. Of the 1042 counties, only 51 had all of the year’s fishing trips originating within the county (black); 148 counties in the study area were not reported as fishing destinations (white).

### Fishing Trips to Multiple Counties

Among the 2576 survey respondents, 68% fished in more than one county, with a maximum of 19 different counties visited by a single individual in the survey year (Fig 4). Counties that hosted anglers that also fished in other counties in that year were widespread across all 12 states in the study area. Of the 894 counties in the study area where fishing was reported, 99% of the counties hosted anglers who also fished in another county in that year.

**Fig 4.**
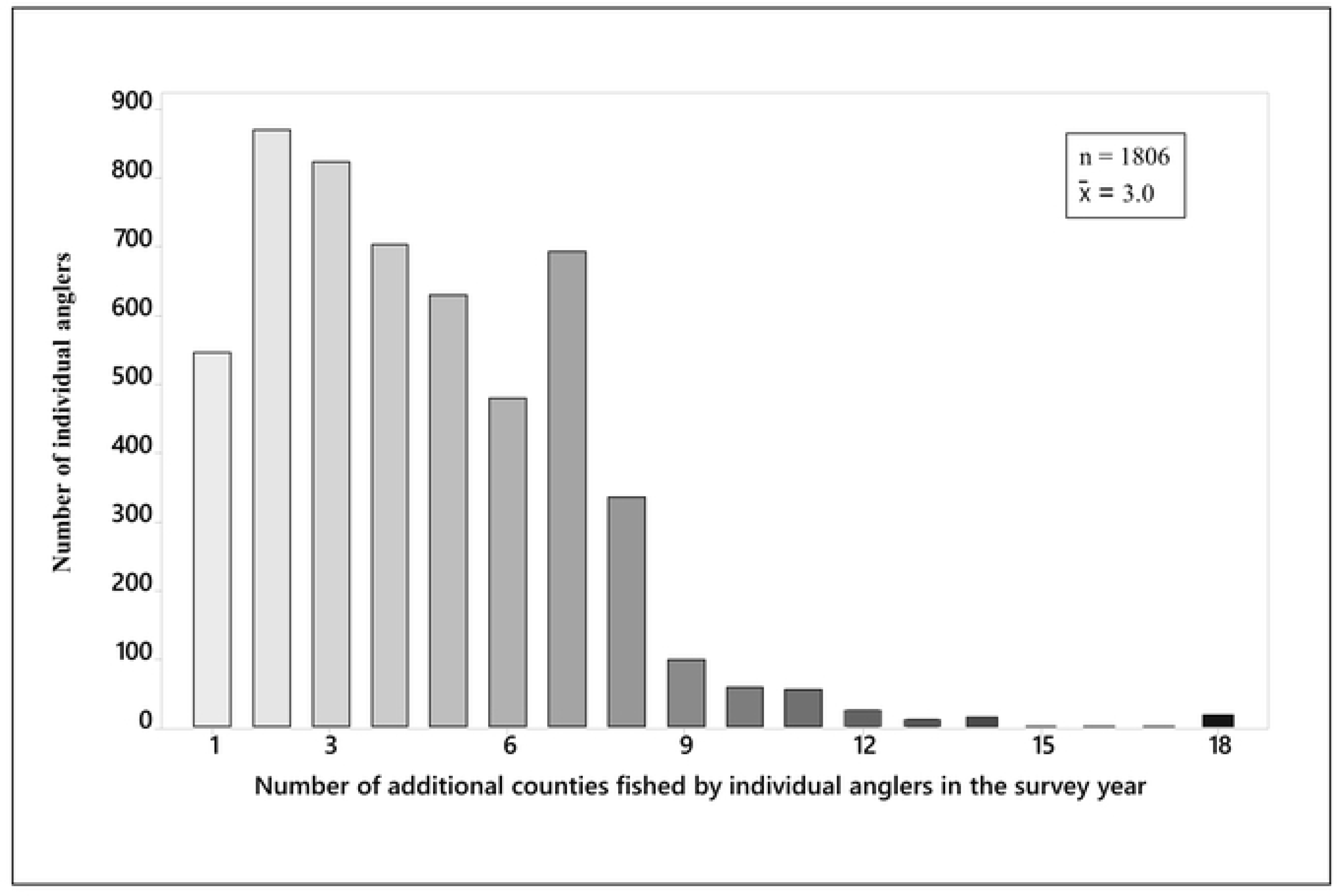
Anglers who fished in multiple counties. Among the 2576 survey respondents in the 13 US states surrounding the Great Lakes and upper Mississippi River basins, 1806 individual anglers (70%) fished in more than one county during the survey year. For those anglers who fished in multiple counties, the mean number of additional counties fished was 3.0. One angler fished in their home county and 18 additional counties during the year.

### Links between counties and between watersheds

Tracking the fishing destinations of the 1756 individual anglers who fished in multiple counties over the study year, we estimated and mapped the weighted sum of all direct linkages and the connecting paths for anglers that moved among counties (Fig 5). The estimated 25.8 million links along 8633 pathways between counties in a single year demonstrate the very high likelihood of spread by anglers and the potential distribution of aquatic invasive species between and among counties across the broader region. Of the 881 counties that hosted anglers who also fished in other counties, the estimated total numbers of direct linkages by anglers between counties ranged from 1199 to 38,957 trips between water bodies in separate counties in a single year. For pairs of counties that were linked by more than 20,000 angler movements, 27 of 29 were connected to major metropolitan areas in the region, such as Buffalo, Cleveland, Chicago, Detroit and Minneapolis-Saint Paul. The highest calculated linkage in the year was between water bodies in Crow Wing and Cass Counties in Minnesota. In addition to areas adjacent to the Great Lakes, the mapped linkages of angler movements emphasized other regions that were highly connected to other counties, such as the inland waters of northern Minnesota, northern Wisconsin, and the Adirondacks in New York.

**Fig 5.**
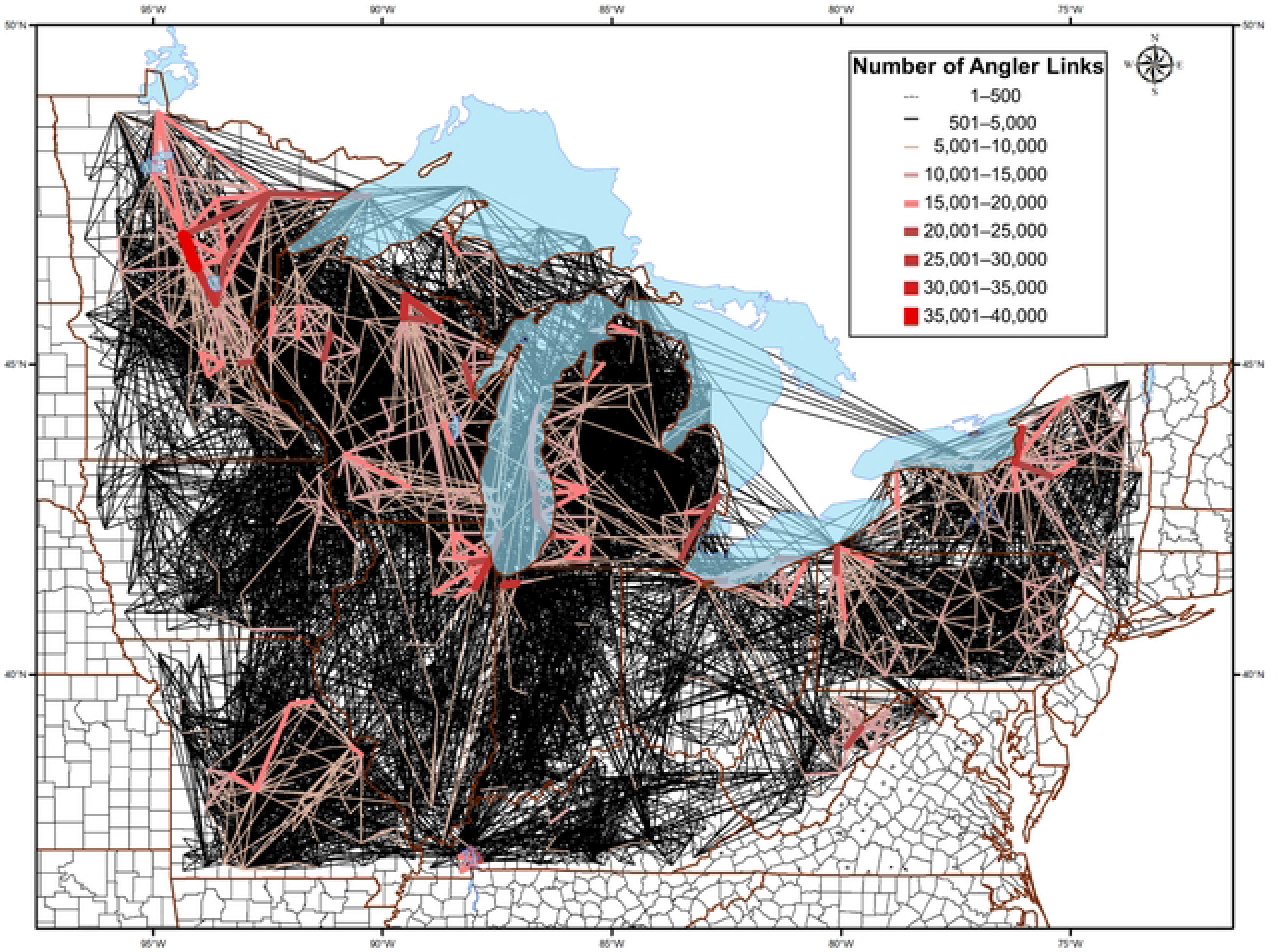
Connections and estimated magnitudes of movements of anglers among counties in the 13 US states surrounding the Great Lakes and upper Mississippi River basins. The estimated 14,766 links among counties in a single year demonstrate the likelihood of spread by anglers and the potential pathways of transfer of aquatic invasive species. Faint black lines represent annual connections between counties numbering from 1–5,000; red lines range from 5,000 (thinner lines), to 40,000 (thick lines) angler trips between counties in a single year.

Because aquatic invasive species invade watersheds rather than counties, we also created a mapped network of connections and movements of anglers among the HU 4 hydrologic units (watersheds) in the broader Great Lakes region (Fig 6). Of the 66 watersheds in the study region, angler movements in the single year connected 61 watersheds along 541 different potential linkage pathways.

**Fig 6.**
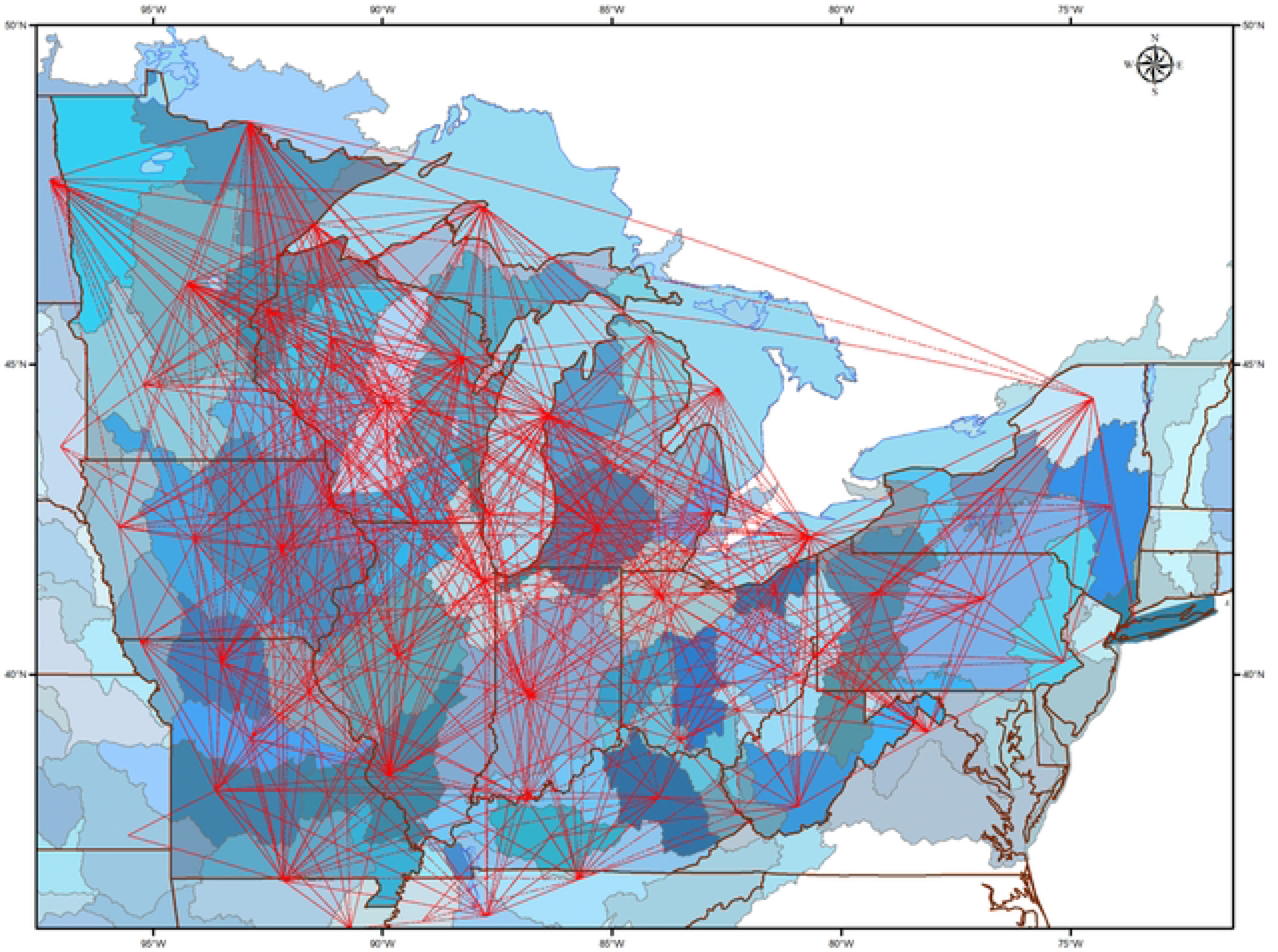
Network of linkages among the 66 watersheds (HUC4 Hydrologic Units) that comprise the 12 US states surrounding the Great Lakes and upper Mississippi River basins. Lines between watersheds represent direct connections made by individual anglers fishing in multiple watersheds in the survey year. The multiple direct connections among watersheds demonstrate the potential for introduction and spread of aquatic invasive species by anglers along linked pathways. The 2576 survey participants were a subset of all anglers fishing in the study region; nevertheless, their fishing behavior connected 92% of the watersheds through inter-basin angler travel along 541 different potential linkage pathways.

## Discussion

We have created an analytical tool, that, in combination with an extensive survey, is both granular and comprehensive in scope, and that provides a stark representation of the challenges associated with attempting to stem the potential transmission and spread of aquatic invasive species by millions of anglers each year. Through the combined efforts of an extensive and geographically widespread survey of thousands of licensed anglers in the Great Lakes region of the US, resulting in a spatially referenced analysis of the behavior of more than 2500 individuals, we estimated that 881 of 1042 counties were linked through 14,700 direct connections of anglers fishing in multiple counties in the study year. Similarly, angler movements directly connected 61 out of the 66 watersheds in the study region, highlighting the interconnected nature and the high likelihood of spreading invasive species among the region’s water bodies.

The study of recreational fishing patterns in inland waters is as complex as it is critical to the management of aquatic invasive species. The scope and inherent difficulties of describing and predicting angler behavior necessitates a data set that is comparable in breadth and magnitude. Our extensive data not only allowed us to generate patterns of connections over a broad geographic area, but enabled us to estimate the behavior of more than 6.5 million anglers with respect to potential of spreading invasive species. In the survey, 68% of anglers fished in more than one county, with one notable individual fishing in 19 different counties in a single year. Counties that hosted anglers that also fished in other counties in that year were widespread across all 12 states in the study area. Of the 894 counties in the study area where fishing was reported, 99% of the counties hosted anglers who also fished in another county in that year.

Through extrapolations, the calculated weighted sums of fishing trips with destinations to each individual counties ranged from 1199, to more than 1.95 million to a single county during that period, with an estimated 143,000 trips to a single county by anglers from another county. Although this was a broad-scale undertaking, it does not include the movements of anglers into the region from the remainder of the US and the provinces of Canada. Therefore, our documentation of the extensive connections and estimated magnitudes of angler movements within the Great Lakes region are likely a considerable underestimate.

The potential for aquatic recreationists to move AIS is of particular concern—and is especially in need of further attention and research--because not all boaters and anglers clean their equipment in ways that will remove AIS. For example, it was estimated that two-thirds of registered boaters in the Great Lakes states of Wisconsin and Michigan do not always clean their boats [34]. A survey revealed that more than half of anglers moving from one body of water to another in six Great Lakes states always removed mud, plants, and fish or animals; nearly three-quarters drained all water-holding compartments; and fewer than half always dried, disinfected, or rinsed their equipment and boats with hot water [10] . In the UK, 64% of anglers surveyed used their equipment in more than one inland catchment within a two-week period, increasing the potential of serving as vectors of AIS [4]. Only 21% of them cleaned and dried their equipment after every use, and only 1% of 446 anglers who fished in other countries and fished at least once every two weeks cleaned their equipment after every use. Even under conditions of extreme desiccation, some invasive species found in Great Lakes waters can survive for several weeks [45,46].

The use of large datasets to generate broad scale patterns of linkages among water bodies can be a powerful tool in studies of AIS. Undoubtedly, managers would benefit from knowing which bodies of water are most at risk. Muirhead and MacIsaac [23] argued that it was important to be able to predict the spread of AIS in spatially explicit ways. By doing so, management efforts can focus on the most susceptible areas and attempt to block movement over the routes that AIS are most likely to travel. To that end, tracing and modeling the movements of millions of individual anglers is the first important step. Using such a tool can also help us move beyond merely establishing physical connections among water bodies. Constructing a weighted network of connections will empower us to focus on specific pathways that are important in at least a few ways: sheer magnitude of connectivity between fishing locations; source areas experiencing an AIS outbreak; vulnerability of previously unexposed destination sites; and areas that are experiencing more than one of these issues. Overall, this tool would not be limited to a single organism or location, but rather can be readily adapted to any species anywhere. Such an analytical approach would be useful for resource managers to target areas at risk, or even common travel routes and to reduce AIS movement by recreational users, such as communication campaigns, boat wash stations and equipment inspections.

Identifying potential routes of transmission can be instrumental in proactive measures and early response efforts to control undesired species introductions. Indeed, the identification and management of pathways to prevent the introduction and establishment of invasive species is a key target in the Convention on Biological Diversity’s Aichi Biodiversity Targets for 2020. An important obstacle to overcome is reducing the unpredictability and overcoming the ubiquity by identifying the most common connections between water bodies linked by movements of anglers. Because many of the modeling and tracing tools are compatible with each other, there is great potential in techniques that employ combinations, such as behavioral data combined with probability models; augmenting eDNA sampling with informatics [12]; or complementing surveys with simple risk models [4] or with more complex likelihood models. For recreational fishing, angler apps, [47], can provide a distributed base of anglers, and citizen science programs, more generally [8], could be used in conjunction with a network model to estimate and predict probabilities of spread of aquatic invasives. Overall, there are several promising approaches that combine models and empirical assessments to tackle such a sprawling and complex set of issues and as the basis for predicting and estimating occurrence, distribution pathways, and likelihood of spread of invasive species among water bodies.

Our blended approach complements much past work that has focused on assessing the risks of AIS transport to and from specific water bodies. In the Great Lakes region, this type of work has been conducted in Illinois [48], Ontario [49,50], Michigan [51], and New York [52]. Part of the challenge with this level of spatial resolution is relying directly on gathering data on the movement of recreationists into each individual location—there are too many water bodies and too many widely distributed recreationists to establish a level of risk for every lake. It is also valuable to look beyond individual bodies of water and focus on the likelihood of AIS transfer between areas of land, such as counties or watersheds that contain multiple bodies of water. Exploring the movement of recreational users between counties, or ecologically meaningful units of land such as watersheds, would provide portraits of risk that may be coarser than an effort focused on individual water bodies, but easier to apply over a large geographic area. This type of analysis would be relevant to current policy questions, such as the concern about AIS transfer between the Great Lakes and Upper Mississippi and Ohio River basins.

Our spatial socioecological analysis, importantly, does not engage the movement of any particular AIS, but looks at potential pathways of transmission. By also expanding the focus of analysis beyond individual AIS, which has been the typical approach, there would be an opportunity to complement past work. Leung et al [51] argued that we should focus on identifying the pathways over which AIS may move, rather than the likelihood of particular AIS arriving in an area, because pathways facilitate the spread of multiple species.

There are varied approaches to monitor, retard, or halt the spread of aquatic invasive species into freshwater environments. In our current study, we quantified and mapped the fishing locations of licensed anglers as a proxy for the potential risk of spreading AIS between counties and watersheds in a 12-state region comprising the Great Lakes and Upper Mississippi and Ohio River basins. We leveraged our survey data on all of the locations fished by thousands of anglers in a single year, created a network of connectivity to identify angler-mediated links among counties and watersheds, and generated weighted estimates for the movements of millions of individual anglers. By combining these tools, we were able to contribute a powerful means to highlight magnitudes and pathways of potential risks of spread of a number of aquatic invasive species across an extensive and diverse region.

## Acknowledgements

We thank Nancy Connelly, Greg Poe, and Richard Ready, who worked with us on the design and implementation of the survey, which generated the data set we analyzed in this study.

